# Fluctuations and freezing of biofilm-air interfaces

**DOI:** 10.1101/2024.05.14.594239

**Authors:** Pablo Bravo, Peter J. Yunker

## Abstract

The study of interfacial roughening is common in physics, from epitaxial growth in the lab to pioneering mathematical descriptions of universality in models of growth processes. These studies led to the identification of a series of general principles. Typically, stochastic growth produces an interface that becomes rougher as the deposit grows larger; this roughening can only be counteracted by mechanisms that act on the top of deposit, such as surface tension or surface diffusion. However, even when relaxation mechanisms are present, interfaces that continue to grow stochastically continue to change; new peaks and troughs emerge and disappear as stochastic growth produces a constantly changing, dynamic interface. These universal phenomena have been observed for bacterial colonies in a variety of contexts. However, previous studies have not characterized the interfacial phenomena at the top surface of a colony, i.e., the colony-air interface, when activity is only present at the bottom surface, i.e., the colony-solid interface, where nutrients are available, over long times. As traditional interfacial roughening models primarily focus on activity occurring at the top surface it is unclear what phenomena to expect over long times. Here, we use white light interferometry to study the roughening of bacterial biofilms, from many different species. We find that these colonies are remarkably smooth, suggesting that a mechanism of interfacial relaxation is at play. However, colonies remain remarkably smooth even after growing large. We discover that topographic fluctuations “freeze” in place, despite the fact that growth continues for hundreds of microns more. With simple simulations, we show that this emergent freezing is due to the dampening of fluctuations from cell growth by the cells between the growing zone and the surface. We find that the displacement field caused by a single perturbation decays exponentially, with a decay length of *δL*. In line with that observation we also show that the topography ceases to change when perturbations are a distance *δL* away from the surface. Thus, over-damped systems in which activity occurs at the bottom surface represent a distinct class of interfacial growth phenomena, capable of producing frozen topographies and remarkably smooth surfaces from spatially and temporally stochastic growth.

## 1 Introduction

Surfaces and interfaces have long been of great interest for their rich physics and complex mathematics[1, 2]. Their characterization led to the discovery of the general rules governing roughening processes and identification of universality classes that emerge accordingly; results of these studies have been used to elucidate physical properties and underlying dynamics of interfacial growth processes[3]. Stochastically growing interfaces typically are self-similar and become rougher as they grow larger[4]. Their only constant is change: topographies continuously vary, as topographic peaks and valleys emerge and disappear over time. However, traditional interfacial roughening features activity at the “top” or outer surface–growth, adhesion, surface tension, diffusion, and more occur at the interface as it grows. It is thus unclear what sort of interfacial phenomena to expect when activity occurs at the bottom surface.

Bacterial colonies, surface-attached communities of microbes encapsulated in extracellular polymeric substances[5, 6], feature two interfaces: one at the bottom where the colony contacts the solid surface, and one at the top where the colony contacts the air above it. Both interfaces feature gradients, for example in nutrients and oxygen; these gradients produce drastically different activity levels at different interfaces[7, 8]. While previous studies discovered that growing bacterial populations exhibit traditional interfacial roughening processes, they always studied the interface where activity, i.e., growth, occurs. For example, multiple groups have characterized the roughening the edge of expanding colonies[9, 10, 11]. Another characterized the air-colony interface over a limited amount of time during which activity occurs throughout the colony[12]. A third observed roughening in a colony growing into porous media [13]. However, as colonies on the air-solid interface continue to grow, reproduction typically occurs in the bulk of the colony, far from the interface, as nutrients diffuse into the colony from the agar gel. Much less is known about what interfacial phenomena occur at the colony-air interface, i.e., where reproduction does not occur, over long periods of time. While the mean dynamics of vertical growth were previously characterized, little is known about the behavior of topographic fluctuations [14]. It is thus unclear if such interfaces are self-similar, if they experience roughening, or if there are any general principles that can be discovered.

Here, we characterize the topographic dynamics of biofilms growing on agar pads at the solid-air interface.

White-light interferometry enables the characterization of the interface with high-resolution (1 nanometer off-plane, 0.174 micrometers in-plane) at a relevant temporal scale (30 minutes). We use openly available interferometry data[15] for colonies of multiple bacterial species(*Aeromonas veronii, Eschericia coli, Kleb-siella pneumoniae, Vibrio cholerae, Bacillus cereus*, and *Staphylococcus aureus*); these data produced one previous publication [14]. This cohort of microbes provides a wide-range in cellular size, shape, EPS production, as well as a wide range of parameters controlling their vertical growth. We find that after an initial period of roughening (as previously reported by [12]), colony topographies smooth and then freeze, even as the colony increases its height by more than a factor of 10. This topographic freezing was unexpected, especially as these colonies are self-similar and exhibit power-law scaling of various quantities across many orders of magnitude. With simple, soft particle-inspired simulations, we show that topographic freezing is the product of the fact that stochastic fluctuations from reproduction occur near the bottom layer of the colony and are then damped by mechanical interactions with particles above. Once the colony is large enough, cells between the active and top layers shield the top from mechanical fluctuations, effectively freezing the topography.

## 2 Results

### A Biofilm topography

The topographic data studied corresponds to measurements across the center of bacterial biofilms (agar-to-agar) for 48 hours after inoculation (Figure 1A). We define the biofilm topography as the height profile of the biofilm-air interface, after removing the long wavelength signal (500 µm) in the lateral direction (Figure 1B). We do this to remove the long-length scale geometry (e.g., [16]), and focus on interfacial fluctuations. Interferometric objectives have slope limits based on their numerical aperture, i.e., they are unable to measure regions that are too steep, as reflected light does not return to the objective. For these reasons, we focus on relatively smooth biofilm morphologies, that is, mature colonies that do not exhibit buckling or wrinkling. Biofilm topographies remain relatively flat; the largest fluctuations remain on the scale of a few micrometers, representing a few cell lengths in amplitude over the 2 mm long profile (Figure 1C).

**Figure 1.**
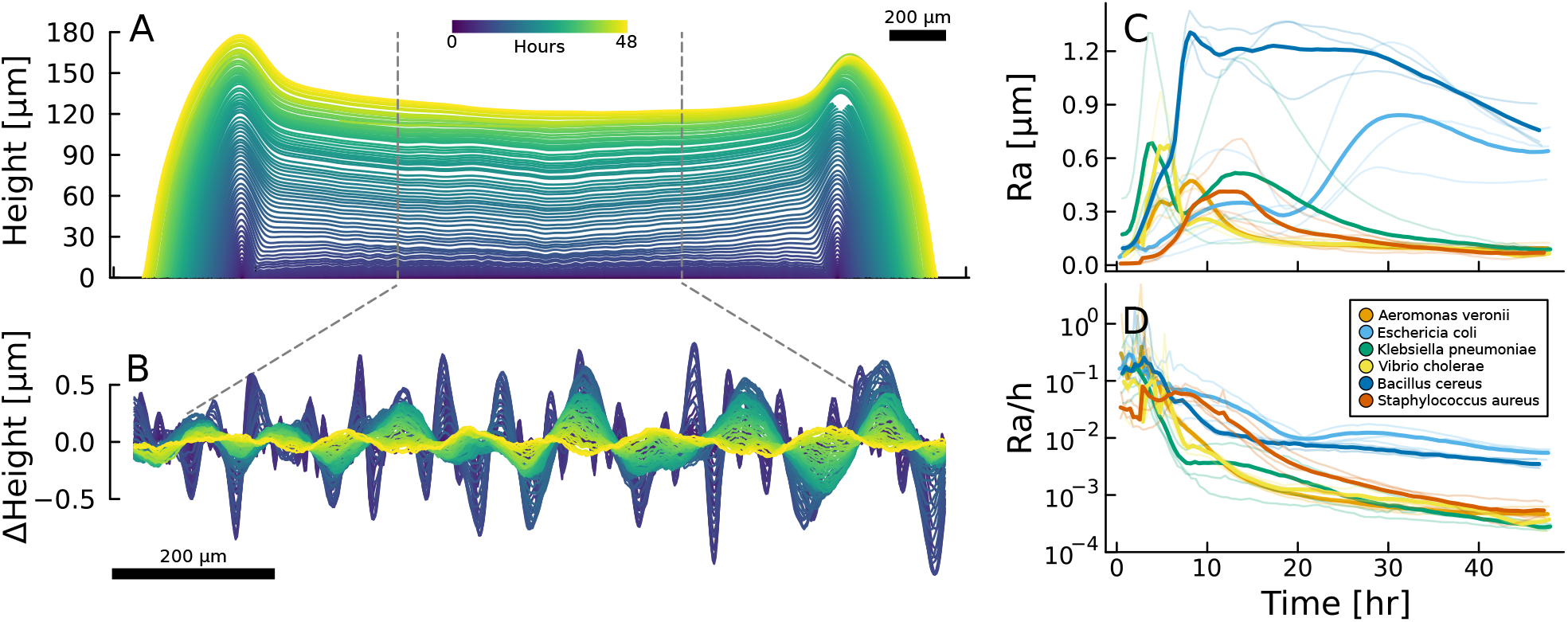
Interface evolution in bacterial topographies. (A) 48-hour interferometric timelapse of a *Staphylococcus Aureus* colony grown in LB-Agar. (B) Fluctuations in the middle 2mm of the colony, a Butterworth Filter with *λ* = 500µm was applied to each individual measurement. (C) Arithmetical Mean Height (Ra) of the substracted profiles, solid lines represent the average of 3 replicates, individual measurements shown in the transparent lines. (D) Normalized fluctuations by their average biofilm height, profiles grow increasingly smoother as they develop, with fluctuations being 3 or 2 orders of magnitude smaller than the colony thickness.

To quantify the characteristic fluctuation amplitude of these surfaces, we calculate the arithmetical mean roughness value 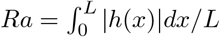, where *h* is the colony height and *x* is the position parallel to the surface. Ra represents the size of the deviations from the mean height of the profile (Figure 1C; for other ISO metrics see Figure S1). The dynamics are characterized by a rapid increase in *Ra*, followed by a subsequent decrease. Two microbes deviate from this norm: *Eschericia coli*, which features two peaks and two decreases, and *Bacillus cereus*, where *Ra* remains relatively high for many hours after the peak before finally decreasing.

To further contextualize the magnitude of the fluctuations in relation to the average height of the biofilms *h*, we calculate *Ra/h*, which represents a unitless metric comparing the characteristic amplitude of fluctuations to the mean biofilm height. Early on, we see that fluctuations correspond to 10% of the total height. As development progresses, this metric decreases to 1% in *E. coli* and *B. cereus*, and 0.1% for *Aeromonas veronii, Klebsiella pneumoniae, Vibrio cholerae*, and *Staphylococcus aureus* (Figure 1D).

Remarkably, these *Ra* values are even smaller than those of polished stainless steel, despite the fact that these colonies experience stochastic growth up to heights equal to hundreds of cells. In fact, *Ra* is less than the size of a single cell. It’s important to consider that the measurements are not obtained through fluorescence, and extracellular matrix products can fill in interstitial regions between cells and at the biofilm-air interface. For comparison, *Ra* for colonies grown into porous media is on the order of 10 of µm after 37 hours of growth [13].

### B Characterization of Smooth Fluctuations

To understand how these air-solid interface biofilms develop into such remarkably smooth interfaces, we next characterize the fluctuations across multiple length scales. By exploring surface topography across multiple lengthscales, one is able to extract more information about the perpendicular and lateral directions. For universal growth processes, many metrics used to characterize the surface morphology, like the interfacial width or power spectral density, exhibit power law scaling with respect to the length scale of observation. This scaling behavior is often related with concepts of fractality, self similarity, and self organized criticality, enabling the identification of the presence or absence of underlying universal growth processes by said exponents.

#### Power Spectral Density

We start by calculating the power spectral density (PSD) *S*(*k,t*), which represents how fluctuations are distributed across different wavenumbers *k*.

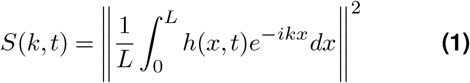

This approach will enable us to identify characteristic length scales in each profile, and, by studying the same interfaces over time, how these length scales change during growth. During the initial growth, colony PSDs primarily exhibit fluctuations between *k* = [2*e* − 3, 1*e* − 2]*µm*^−1^; these length scales likely emerge from the fact that cells are inoculated at a relatively low concentration, and are separated from each other; the topographic initially represents a combination of cell surfaces and the agar-air interface. As colonies mature, after 10-15 hours, the PSD begins to exhibit power-law-like scaling across all the sampled wavenumbers in the form *S*(*k, t*) ∼ *k*^−*ν*^, with *ν* ≈ 2.6(Figure 2A). This power-law behavior extends across 3 orders of magnitude(*k* = [1*e* − 3, 1*e*0]µm^−1^). This measurement corroborates previous characterizations of the PSD in 3-dimensional hydrogels[13], with a similar exponent of *ν* ∼ 2.3. This power-law scaling, and thus the absence of a characteristic wavelength in the PSD suggests that, despite their smoothness, these biofilm-air interfaces exhibit characteristics of self-similar systems.

**Figure 2.**
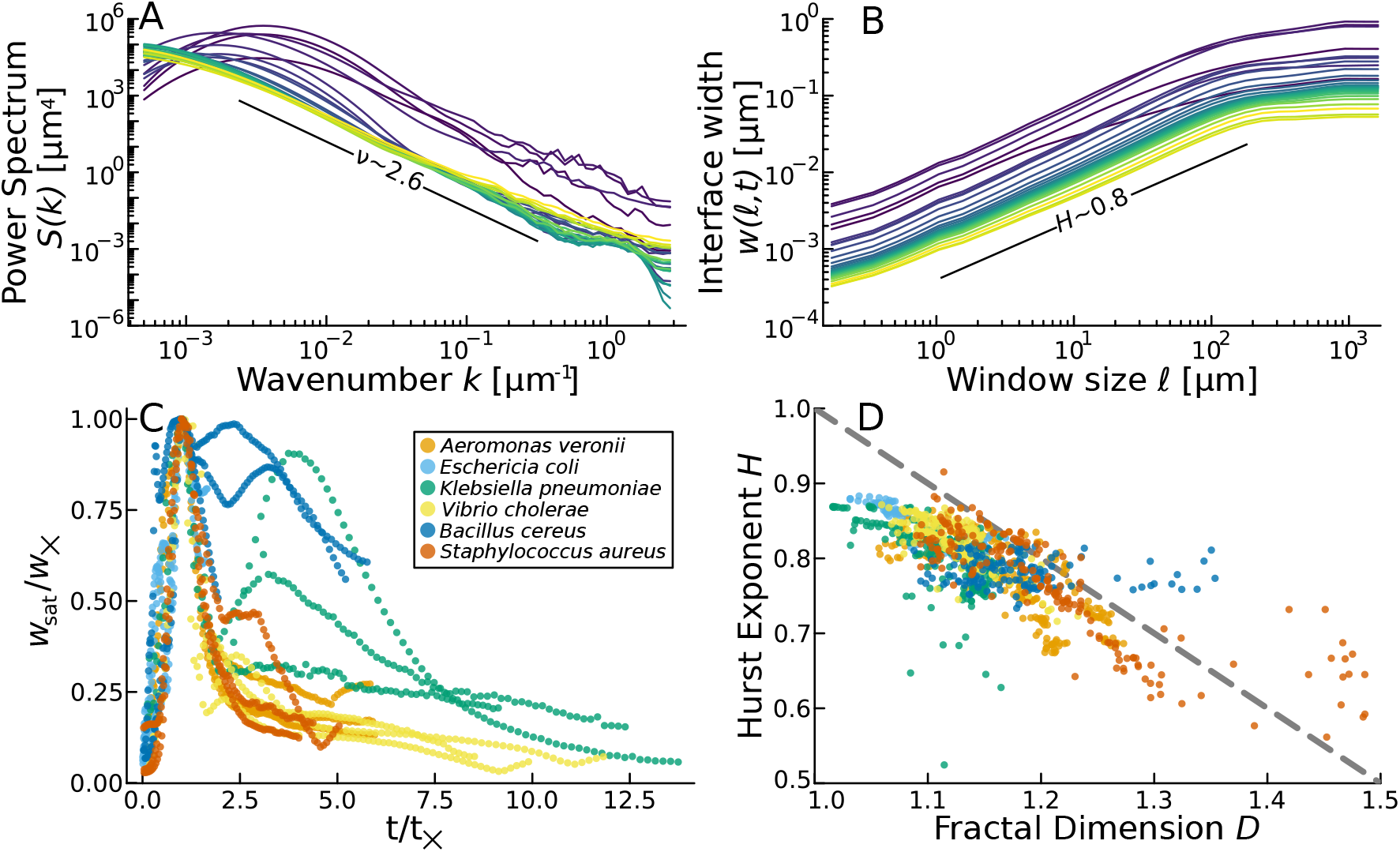
Scaling characterization of biofilm-air interfaces. (A) Power Spectral density of *Vibrio cholerae* biofilm colonies growtn in LB-Agar. As the colony develops, the PSD exhibits scaling behaviour with an exponent *ν* ∼ 2.6. (B) Interface widths of the same colony, the spatial scaling corresponding to the roughness remains stable at *H* ∼ 0.8, while the saturation width exhibits an initial increase followed by a decrease over time.(C) Normalized dynamical scaling behavior of the cohort of bacteria measured. Each timelapse was individually normalized in relation to the maximum interface width *w*_*×*_ and its respective time *t*_*×*_, in a traditional system exhibiting dynamic scaling, the values would remain at a constant saturation width after reaching the transition regime. (D) Phase-space for the Mandelbrot self-similarity criteria 2, marked as the gray dashed line. Only points after the transition time *t*_*×*_ is shown, thus highlighting the conflicting between dynamic scaling and self similarity behaviors of the biofilm-air interfaces.

#### Self-similarity

The characterization of surfaces and interfaces has long featured the search for self similarity: the statistical invariance in which the height profile *h*(*x*) scales as *λ*^*H*^ when multiplying *x* by λ. The general significance of this concept is that the object looks roughly the same at any scale. To first investigate if these interfaces are self-similar, we plot *h*(*x*) profiles on different scales. Indeed, we observe qualitatively similar fluctuations across 4 orders of magnitude(Figure S2). We quantitatively test if these interfaces are in fact self-similar. To do so, we utilize Mandelbrot’s criteria to test for self-similarity[17]. This approach involves characterizing the structure over asymptotically large length scales with their roughness/Hurst exponent *H*, as well as asymptotically short length scales through their Fractal Dimension *FD*, and then checking if *H* and *FD* obey the following relation [18]:

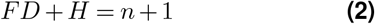

where *n* corresponds to the “base” dimension (for a 1-dimensional profile *n* = 1, for a topographic map *n* = 2). To calculate *H*, we first need to calculate the local interface width *w*(𝓁, *t*), a metric describing the standard deviation in height calculated over sub-sections of different sizes, 𝓁, in the horizontal axis:

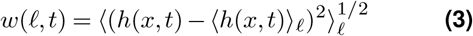

where *h* is the height profile, *x* the horizontal direction, and t the elapsed time since inoculation. The local interface width (Figure 2B) can be represented using three scalar metrics that capture the rough behavior of the function, defined by a scaling regime and a saturation regime for a given profile: *w*_*sat*_, the saturation width, identical to the standard deviation of the full profile, 𝓁_×_, the length scale at which the profile reaches the saturation width, and the Hurst exponent H, the scaling exponent for the roughness exponent at small 𝓁, which follows the power-law: *w*_𝓁_ ∝ 𝓁^*H*^ for 𝓁 < 𝓁_×_.

After an initial period of steadily increasing roughness (0-8 hours), we observe that the profiles reach a steady state around *H* ∼ 0.8, across over 2 orders of magnitude in both 𝓁 and *w*(𝓁, t) (Figure 2). Power-law scaling and roughness exponents have previously been measured for biofilms: in the context of expanding front across an agar plate *H* ∼ 0.78[19] and *H* ∼ 0.67[11], for 3-dimensional colonies growing in porous media *H* ∼ 0.7[13], and the biofilm-air interface at early times *H* ∼ 0.6[12]. White-light interferometry measurements presented in this work expand on this previous literature; we find excellent power-law behavior over nearly three orders of magnitude fitted(𝓁 = [1e − 1, 1e2] µm). We next seek to calculate the fractal dimension *FD*. This metric is a statistical index of the complexity of the profile, as observed at different length scales. After standardization of each profile along both axes, we utilize the correlation-sum method[20] detailed in Datseris *et al*.[21] to estimate the fractal dimension. We observe that at early times, values cluster around *FD* ∼ 1.5 for all colonies: this is expected since during the first measurement the structure of the interface is mostly determined by the inoculation, and not the developmental process. *FD* then steadily decreases over the first 10 hours, until values reach *FD* ∼ 1.2 ± 0.1(Figure S5).

We now have everything necessary to determine if these remarkably smooth interfaces are in fact self-similar using Eq. (2). To do so, we plot *H* versus *FD* for all strains and replicates measured. We find that individual time points are clustered close to the *H* +*FD* = 2 line. As these interfaces are, at least, very nearly self-similar, do they grow via interfacial roughening processes, like those seen in [13, 12]?

#### Dynamic scaling

For interfacial roughening processes, the interface width *w* scales with time *t*, in addition to the sampling length 𝓁. It is expected that, before reaching a transition time *t*_×_, the temporal component governs the dynamics as *w*_sat_(*t*) ∼ *t*^*β*^, and after the transition time *t*_×_ the saturation width remains a constant. However, while we observe spatial scaling characterized by the Hurst exponent, we do not observe temporal scaling in biofilm-air interfaces. The saturation width initially increases, but then decreases substantially, even while we see enough activity to increase the average colony height by a factor of 5 over this period (Figure 1A).

We observe different saturation widths as well as transition times for different strains, with replicates exhibiting similar dynamic behaviors. Normalized scaling, both through *w*_sat_ and *t*_×_ show that biofilm topographies do not follow dynamic scaling in the measured 48-hours of growth, and thus they do not exhibit traditional interfacial roughening during this time. In particular, we observe that although *w*_sat_ increases in the first few hours of biofilm development, it then decreases over time(Figure 2C), similar to our previous observation that surface topography becomes less rough on long time scales. It is clear from these results that the characterization of biofilm topographies, even relatively smooth biofilm topographies, is a complex task. We observe some similarities across the microbial cohort we measured, such as the Hurst exponent and the Fractal Dimension *FD*, while also observing very different dynamics in the saturation width *w*_sat_. However, we consistently observed another phenomenon of interest across all samples: topographic *freezing*.

### C Freezing of the biofilm topography

The temporal evolution of biofilm topography has two stages during growth: (i) active and (ii) frozen (Figure 3A). We call the time at which colonies transition from active to frozen *t*_×_. The active regime shows some variation from strain-to-strain, but the frozen regime is consistently observed in all of our experiments (Figure S3). During the active regime, topographies rapidly change; during the frozen regime, topographies appear not to change, or to solely decay in amplitude at a slow rate.

**Figure 3.**
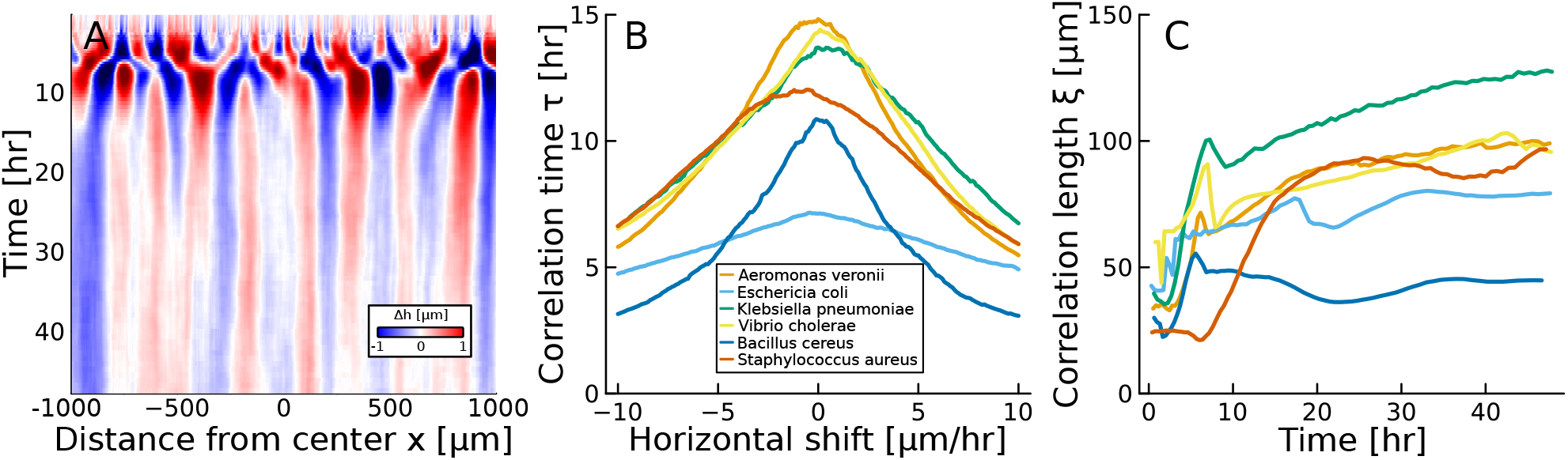
Topographic freezing of biofilm-air interfaces. (A) Kymograph showing the biofilm-air interface for a colony of *Aeromonas veronii*, peaks are colored red (+1*µm*) and valleys in blue (−1*µm*). The freezing of the interface is clear after t = 10 hours. (B) Temporal correlation times after the settling of the topography, a total of 100 horizontal shifts were sampled for each timelapse. Only the mean value between the 3 same-species replicates is shown, the standard deviation roughly corresponds to 1*/*4 of the mean. Highest correlation times around 0 mean that there is no travelling waves in the topographies. (C) Correlation lengths for surface topographies, after colonies settle, there is a slow increase in correlation length *ξ*, implying the presence of a diffusion-like relaxation force.

As an alternative to being frozen, fluctuations could be moving across the surface as a travelling wave. To quantify if travelling waves are present, we calculate the correlation times τ across the profile in this regime, at different horizontal shifts. This metric, when averaged over all positions, would capture the speed of a travelling wave if one was present. However, for all samples, we observe that the highest correlation time is at τ = 0, with correlation times ranging from 6 to 15 hours(Figure 3B). This indicates that the topographic features are not propagating laterally, but rather remain stationary or “frozen” in place.

We next look at how characteristic length scales change in the frozen regime. To do so, we quantify the correlation length ξ, the average horizontal distance at which a fluctuation decays to 1/*e*. After growth from their initial low density conditions, colonies exhibit a sharp jump in their correlation length, which increases by more than a factor of 2. In the topographic freezing regime, ξ increases steadily with time for most colonies(Figure 3C). While the relationship between ξ and time is not perfectly monotonic, the slow increase in ξ with time is roughly consistent with ξ ∼ *t*^¼^ (Figure S4), a phenomenon predicted for wrinkling films [22]. This result, coupled with other metrics linking to the amplitude of these fluctuations(Figure 1C, D) suggest that there is a diffusion-like relaxation force playing a role at the biofilm-air interface.

Topographic freezing represents a striking departure from traditional interfacial roughening behavior. Though colony surfaces exhibit self-similarity, with well-defined fractal dimensions and Hurst exponents, they are unable to follow dynamic scaling relations as they freeze. Why does the topography freeze, despite continued growth of the colony? One possible explanation is that as the biofilms grow thicker, the actively growing region near the agar substrate becomes increasingly separated from the air interface. Stochastic fluctuations in growth at the base of the biofilm may be dampened by the intervening layers of cells before they can propagate to the surface. This would result in a “frozen” topography at the biofilm-air interface, even as the biofilm continues to increase in overall thickness.

### D Topographic freezing in a simple model system

To explore the phenomenon of biofilm-air interfaces freezing, we developed a simple agent-based model that captures the essential features of biofilm growth and mechanical interactions. In this model, cells of different sizes are represented as soft disks growing in a 2D plane, with a solid substrate at the bottom, representing the agar, a free surface at the top, and periodic boundaries in the horizontal direction. Mechanical interactions between cells are modeled using a soft repulsive potential, which allows for local rearrangements and deformations in response to growth-induced stresses (see Methods).

Using this model, we investigate the effect of a single stochastic “growth” event on the surface topography of biofilms. We hypothesize that the main process leading to the change in the biofilm-air interface is cellular reproduction, and thus focus on perturbing the interface by adding a new cell to the colony. Considering that carbon sources are supplied from the biofilm-agar interface, we produce a perturbation in the bottom layer of the colony. In small colonies, the cascade of rearrangements due to mechanical interactions can travel through the whole colony, enabling a single perturbation to change the topography(Figure 4A). On the other hand, in a large colony rearrangements due to the perturbation are damped and distributed across many neighboring cells, with negligible effect on the cells at the top surface(Figure 4B). The topography change is even more extreme when instead of a single perturbation, we iteratively add more perturbations. In the small colony regime only some of the original features remain after the perturbations, whereas in the large colony regime the profile is effectively the same, only displaced to account for the new volume added to the system (Figure 4A,B).

**Figure 4.**
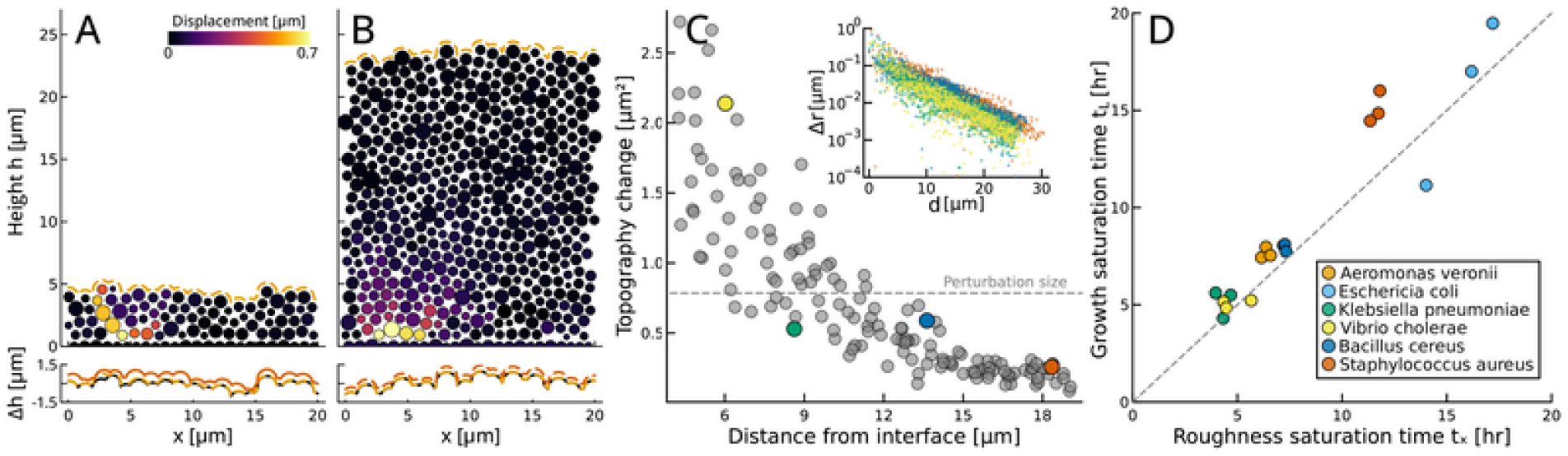
Model for interface freezing. (A) A small (*N* = 100, ⟨*h*⟩ = 4.5*µm, x* = 20*µm*) and a (B) large (*N* = 500, ⟨*h*⟩ = 23*µm, x* = 20*µm*) colonies after a single perturbation in the bottom layer, the colorbar corresponds to the total displacement respective to their original positions after relaxation. In the bottom panels the initial topographies (black) are shown next to the states after 1 perturbation (dashed), and 10 perturbations (solid). (C) Change in the topography given perturbation distance in a large simulation *N* = 1000, ⟨*h*⟩ = 19*µm, x* = 50*µm*. (Inset) Cell displacement Δ*r* in relation to the distance to the perturbation follows exponential decay, colors correspond to perturbation events in the main plot. (D) Correlation between the timing to reach the saturation time *L*, as defined in **??**, and the first “peak” in saturation width *w*_sat_.

To quantify these effects, we explored the behavior of a large simulated colony (*N* = 1000, ⟨*h*⟩ = 19µ*m, x* = 50µ*m*) and observed the change in topography after a perturbation. Recapitulating the aforementioned behavior, perturbations near the interface have a much larger effect on the topography; the amount the topography changes decays with distance from the interface, with the effect on the topography being less than the size of the cell added at a distance of ∼ 10 µm (Figure 4C). We also observe that the variance of the measured topography change decreases with increasing *d*, likely due to finite size effects.

To understand what sets the size a colony must grow to for the topography to freeze, we measured the displacement, Δ*r*, of every cell after a perturbation in a large colony. We found that Δ*r* decreases exponentially with increasing distance from the perturbation, *d*, i.e,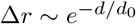 (Figure 4C inset, **??**), where *d*_0_ = 13.33 µm. Thus, the height our model colony had to grow to for the topography to freeze is the characteristic length scale of impact from a perturbation.

These simulations suggest that once the system reaches a characteristic height, perturbations from new cells are substantially damped, preventing fluctuations from modifying the topography. To test this idea with our experimental data, we compare our previous measurements characterizing the times at which colonies reach *h* = *L*[14], i.e., the point after which colony growth rate decreases with increasing height until they eventually reach an equilibrium steady state height. These timings correlate well (*R*^2^ = 0.91) with the timing at which the biofilm-air interface reaches the first “peak” in the saturation width *w*_sat_(Figure 4D), i.e., the point at which roughness ceases to increase. Thus, our experiments are consistent with the idea that topography freezes due to dampening from inactive cells.

## 3 Discussion

In this work, we characterized a diverse cohort of smooth biofilms with white-light interferometry and explored how topographic fluctuations correlate with biophysical growth processes in the colony. Interferometry enabled us to investigate the scaling of fluctuations over three orders of magnitude. We found that after an initial period of increasing roughness, colony topographies freeze, even as the biofilm continues to grow. This frozen state is characterized by long correlation times and steadily increasing correlation lengths, indicating a suppression of lateral feature propagation and a stabilization of topographic structures. The observation of this freezing phenomenon across diverse bacterial species suggests that it arises due to a universal feature of biofilm development. Using a simple mechanical model, we show that the amount cells are displaced by the emergence of a single new cell decays exponentially with distance from the new cell; as reproduction occurs near the substrate, once biofilms grow taller than this region of influence, reproduction no longer affects topography. Finally, we corroborate this model by establishing a temporal correlation between the freezing of the biofilm-air interface with their growth dynamics, highlighting the coupling between mechanical interactions and developmental processes.

In recent years, active interfaces have been studied more frequently, from the interfacial mixing of actin components, to the swarming of bacteria colonizing their environment. Surprisingly, these complex biological phenomena often map well onto previously identified interfacial roughening universality classes. In active matter systems, agents interact with each other and consume energy, leading to rich collective dynamics. While the activity in traditional active matter comes from the locomotion of individual agents, activity can also come from growth, reproduction, and death. These behaviors have been studied in the context of cell tissues, bacterial colonies, and corals, uniting these immotile systems, in part, with traditional forms of active matter. However, the complex biology behind growth, reproduction, and death have limited our ability to elucidate the underlying biophysics behind these diverse systems. Here, we show that topographic freezing occurs for a wide range of bacteria, suggesting that it may be a common phenomenon when growth and reproduction primarily happen far from the interface.

We found that the biofilm-air interface of smooth colonies is self-similar. From a macroscopic view these colonies appear remarkably smooth; however, the interface is complex and irregular across multiple orders of magnitude. From a reductionist perspective, the self-organization in a biofilm colony is largely a product of nutrient gradients and mechanical interactions; thus, simple rules shared across the cohort of microbes we study may be expected to produce similar fractal dimensions and Hurst exponents in the self-similarity phase-space[23](Figure 2D). However, other microbes not studied here may have substantially different underlying rules. Colonies that exhibit wrinkling due to mechanical stresses will differ, as well as colonies that can bypass the nutrient limitations through water channels. Further, microbes that are obligate aerobes will likely reproduce in the middle of a colony, where oxygen and nutrients may both be available. Nonetheless, the conditions necessary for freezing—reproduction occuring far from the air-biofilm interface—likely occur in many systems, inside the lab and in nature.

Scale-free phenomena can also be observed in the power-law scaling of the biofilm PSD, which is maintained even as the overall surface roughness decreases during the later stages of colony development. This fact indicates that the relative distribution of topographic fluctuations remains constant, even as their absolute amplitude diminishes. Such scale-invariant behavior is a hallmark of self-organized criticality and has been observed in a variety of physical and biological systems, from the dynamics of earthquakes to the patterns of neural activity[24].

The roughening and fractal-like behavior has been characterized in biofilms in multiple settings. Early work characterized the fractal patterns in horizontal range expansion and how growth depends on nutrient availability and agar concentration[11, 9, 10]. Characterizations of the morphology in other planes, like growth in 3-dimensional hydrogels[13] and quasi-2D colonies in agar[12], suggest that this self-similarity is an universal behavior in bacterial colonies. In this work we exploit the spatial resolution of interferometry to enable the characterization of power-law behavior across 3 orders of magnitude in 1-dimensional topography profiles. Collectively, these results are characterized by values for the surface roughness higher than what would be expected from random interface growth[25, 26]. Biofilms exhibit large correlations across long scales, and thus suggest that non-local effects couple the interface across multiple scales[11, 27]. Defining the mechanisms behind, and consequences of, these effects, like mechanical interactions, growth gradients, and cell-cell signaling, is an intriguing avenue for future research.

A simple mechanics-based model provides a simple but powerful framework for understanding the complex interplay between growth, mechanics, and surface morphology in bacterial biofilms. This model reproduces the topographic freezing behavior to mechanic interactions due to cellular growth. The spatial heterogeneity in growth activity plays a crucial role in the biofilm-air interface dynamics. If cells grow near the interface then the topography will keep changing; if cells grow far from the interface, the topography will freeze. This model captures the mechanisms underlying topographic freezing; however, a more complete model would be needed to completely capture the dynamics observed in experiments. Incorporating nutrient gradients, as well as stochastic rates of cellular growth, lysis, and reproduction would elucidate how these phenomena interact to produce the complex, dynamic biofilm topography. Computational studies that incorporate these effects have been previously developed[28, 29, 30], but the simulation of large colonies over long timescales is difficult due to the need for immense computational power and memory.

In models of out-of-equilibrium systems, the energy uptake that drives interfacial fluctuations typically occurs either throughout the entire system or near the interface itself. However, for biofilm-air interfaces, the fluctuations arise from stochastic cellular growth and division processes occurring far from the measured interface, at the base of the biofilm. This unique feature has important implications for our understanding of interfacial dynamics in complex biological systems. This work highlights the importance of considering long-range interactions and the role of the system’s internal structure in shaping its surface properties. Furthermore, the observation that these distant fluctuations can lead to a “frozen” topography, rather than the constantly evolving surfaces seen in traditional active matter systems, suggests that the interplay between local growth processes and global mechanical constraints can give rise to novel, non-equilibrium steady states.

Local growth and nutrient gradients are known to change the morphology of bacterial interfaces that contact a source of nutrients[13, 31, 30]. Fingering instabilities can form, enabling cells that are more immersed in the substrate to grow even faster. This phenomenon is counteracted by cell-cell adhesion, acting like a pressure to homogenize the interface[29]. For simple biofilm-air interfaces, the source of this fingering instability is not present, enabling the pressure-like relaxation force to dissipate the fluctuations in the surface topography.

In conclusion, the biofilm-air interface is a rich, under-explored source of information about the biological and physical phenomena occurring beneath the surface. The results presented here provide a foundation for further experimental and theoretical studies aimed at elucidating the physical principles governing biofilm development and function. For example, if the topography freezes but the colony keeps growing, that implies reproduction occurs far from the air-biofilm interface; in contrast, if the interface continues to change, reproduction must be occurring close to the biofilm-air interface. Future work may enable us to connect biofilm topography to more biological and physical processes proceeding deep inside biofilms.

## Methods and Materials

### A Analysis of interferometry profiles

We used the collection of interferometry measurements in the Dryad Repository. We define the topography as the middle 2mm region in the colony, long-range wavelengths(which are products of the agar surface, and the curvature from the coffee ring) were removed by passing topographies through a Butterworth filter with a cut-off of λ = 500 µm.

### B Agent Based Simulation

The system consists of 2-dimensional circular particles, interacting through the potential U

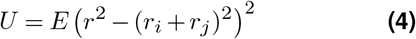

Where *E* is the elastic modulus, *r* the distance between the two particles, *r*_*i*_ and *r*_*j*_ the radii of each particle. For the sake of simplicity we maintained an elastic modules *E* = 100*kPa*. The system consists of periodic boundary conditions in the x-direction, with a solid boundary condition on the bottom and an open boundary condition on the top. In order to set the initial configuration *u*_0_, we initialize *N* cells of radius *r* (*r* ∼ *N*(µ = 0.5, σ^2^ = 0.05)) in a constrained box (φ ≈ 0.9). Then, the simulation is left to relax until the average net force f applied to a cell from its neighbors is below 10−3.

In order to perform a perturbation in the system, a particle of radius *r*_*p*_ is introduced at a position 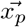. The system is updated until it reaches the same relaxation condition of ⟨*f*⟩ < 10−3. While doing this we keep track of the forces and displacements of each particle during relaxation.

Additionally, for each configuration *u*, we calculate the topography 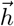. To do this we implement a reference grid of points overlaying the system. For each reference point, the distance to each particle center is calculated. For all values of *x*, the highest value of *y* that is in the neighborhood of a cell, plus an added padding (0.5µ*m*) is returned as the biofilm-air interface(Figure S6).

## Data availability

Interferometry data can be found in the Dryad Repository. Analysis, simulation code, and processed data will be available on Github.

## Supplementary Information

**Figure S1.**
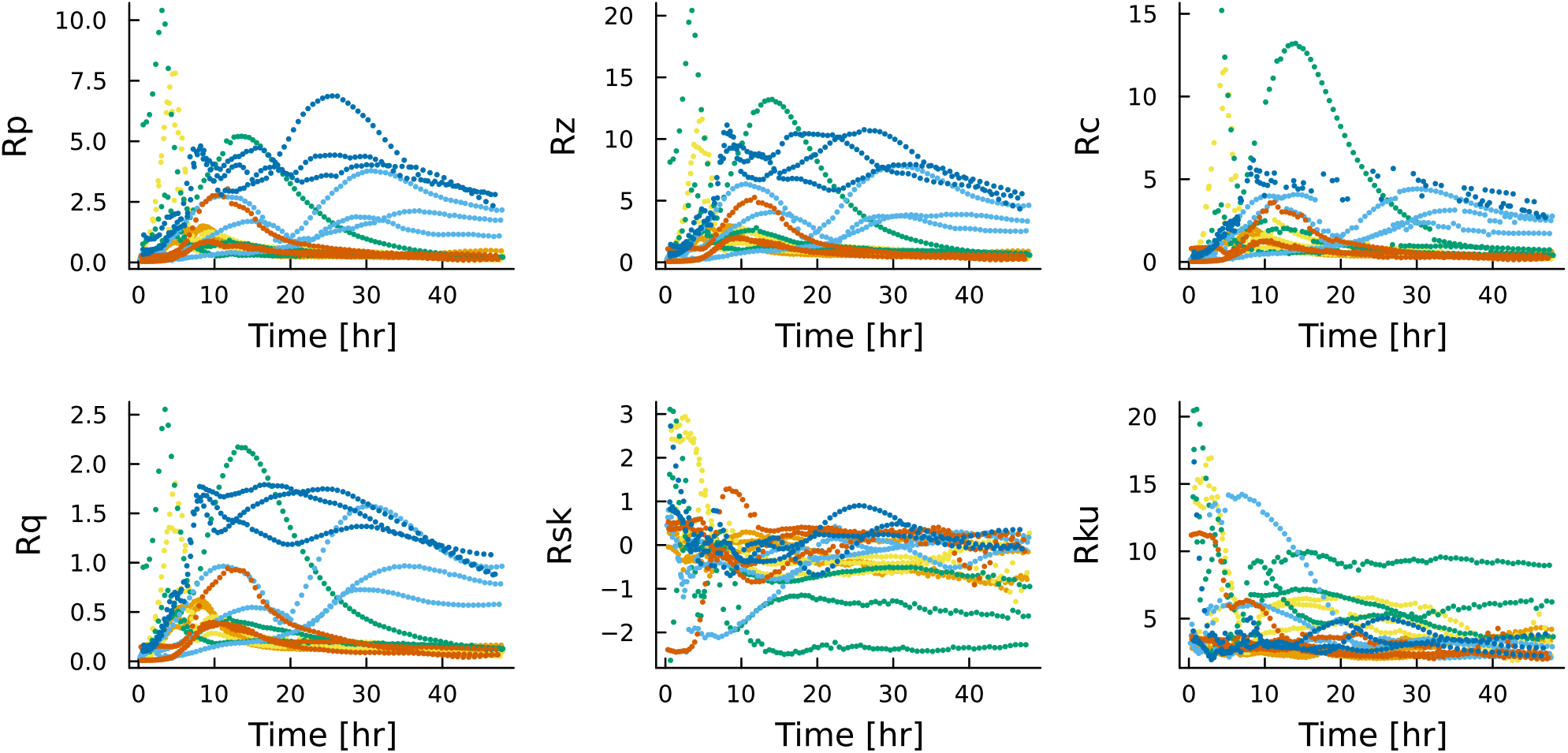
ISO Metrics for biofilm-air interfaces. All calculations are following ISO-4287: Geometrical Product Specifications. Maximum peak height *Rp*, maximum height of the profile *Rz*, mean height of profile elements *Rc*, root mean squared deviation *Rq*, skewness *Rsk*, and kurtosis *Rku*.

**Figure S2.**
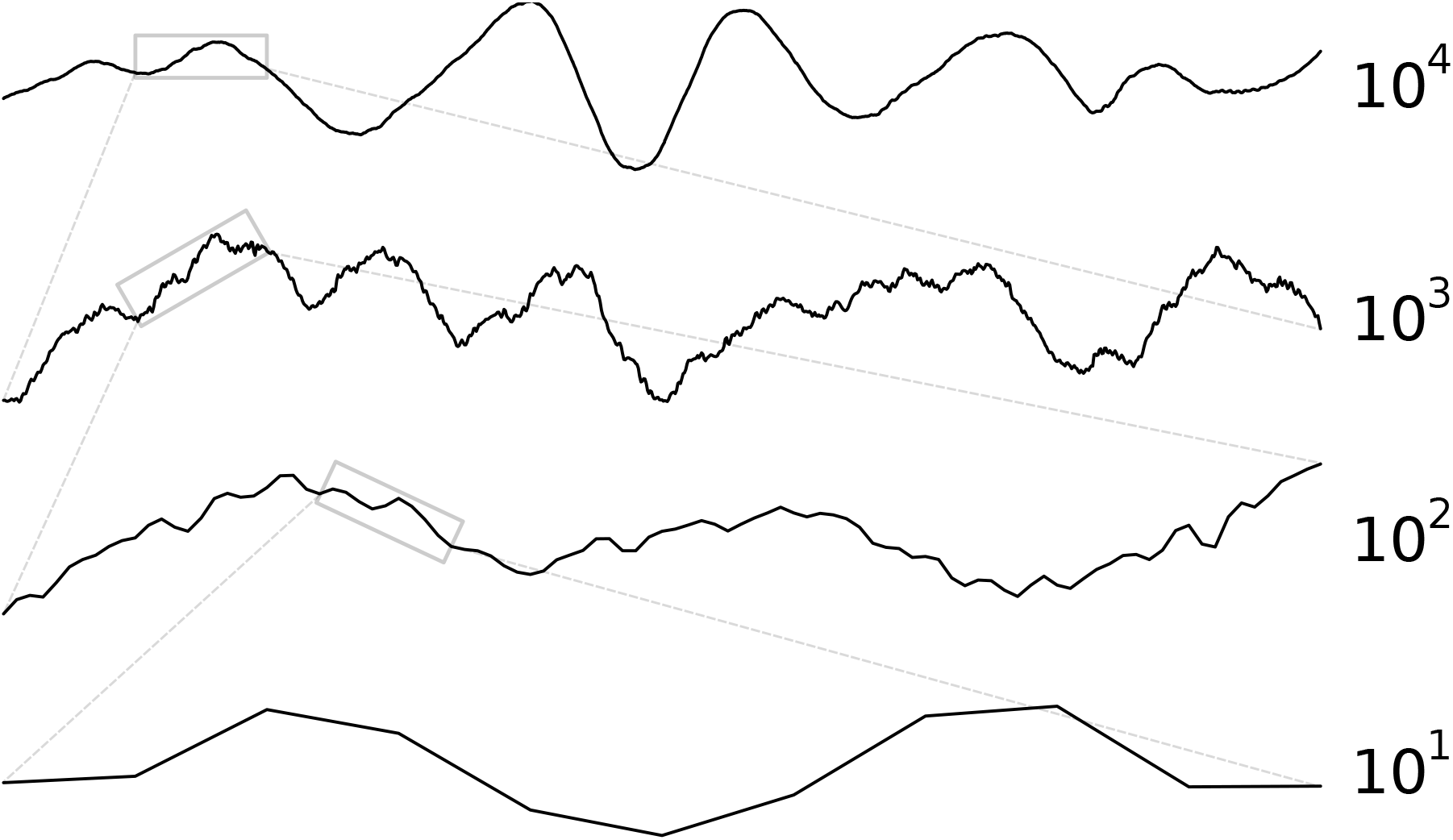
Visual representation of self-similarity in biofilm topographies. The biofilm-air interface of *Vibrio cholerae* at 10 hours of growth, near the transition to “freezing”. Initial slice corresponds to 10^4^ pixels, roughly *L* = 1700*µm*. Each subsection then goes through a Butterworth filter with a cutoff wavelength of *λ* = *L/*2.

**Figure S3.**
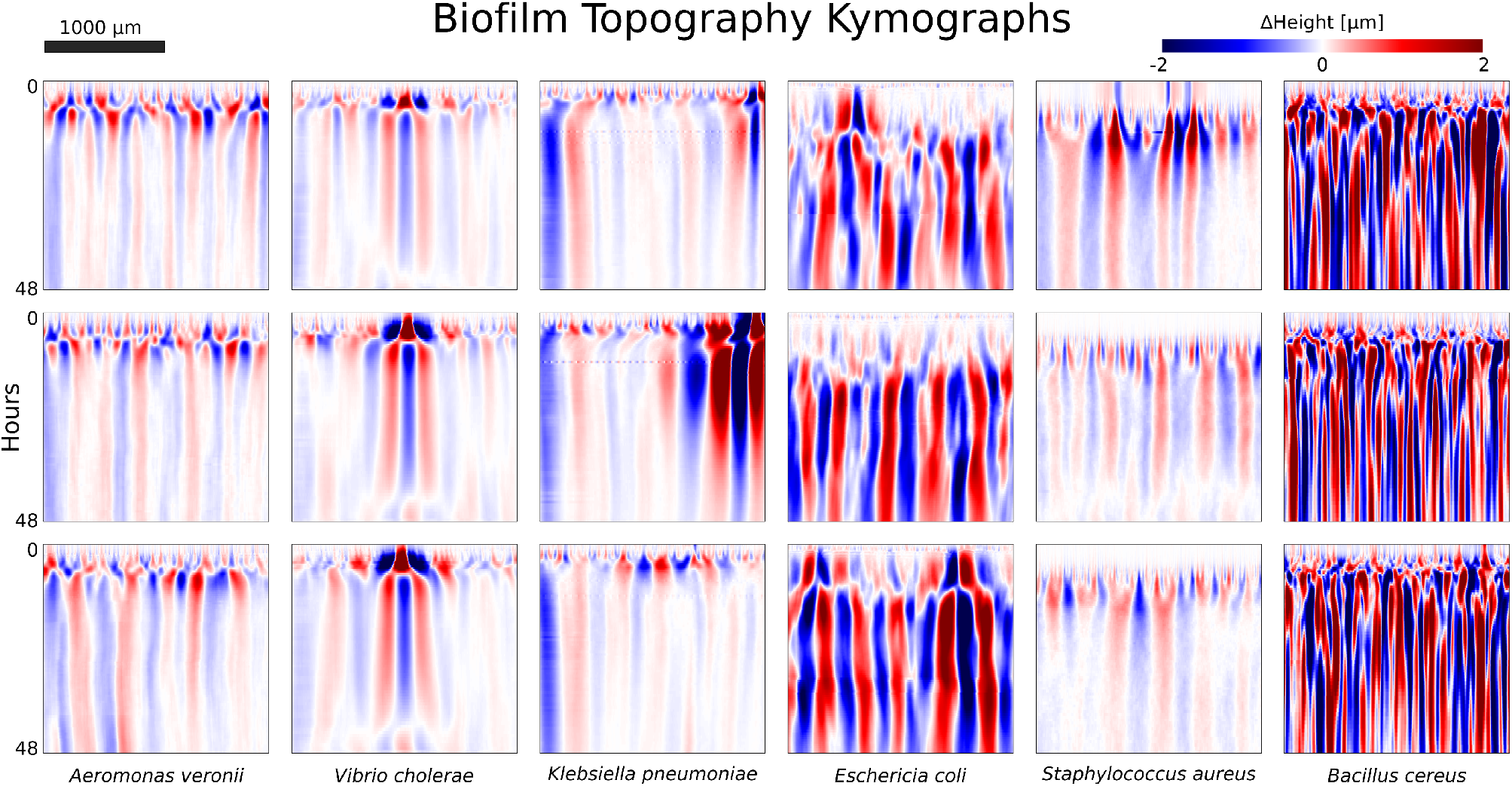
Biofilm-air interface kymographs. Kymograph showing the biofilm-air interface for colonies of *A. veronii, V. cholerae, K. pneumoniae, E. coli, S*.*aureus*, and *B. cereus*. Each species consists of 3 replicates, A (top), B (middle), and C (bottom). All kymographs are using the same diverging colormap: peaks are colored red (+2*µm*) and valleys in blue (−2*µm*). Even if the correlation length varies (width of peaks and valleys), the persistence of the topography after it settles is clear.

**Figure S4.**
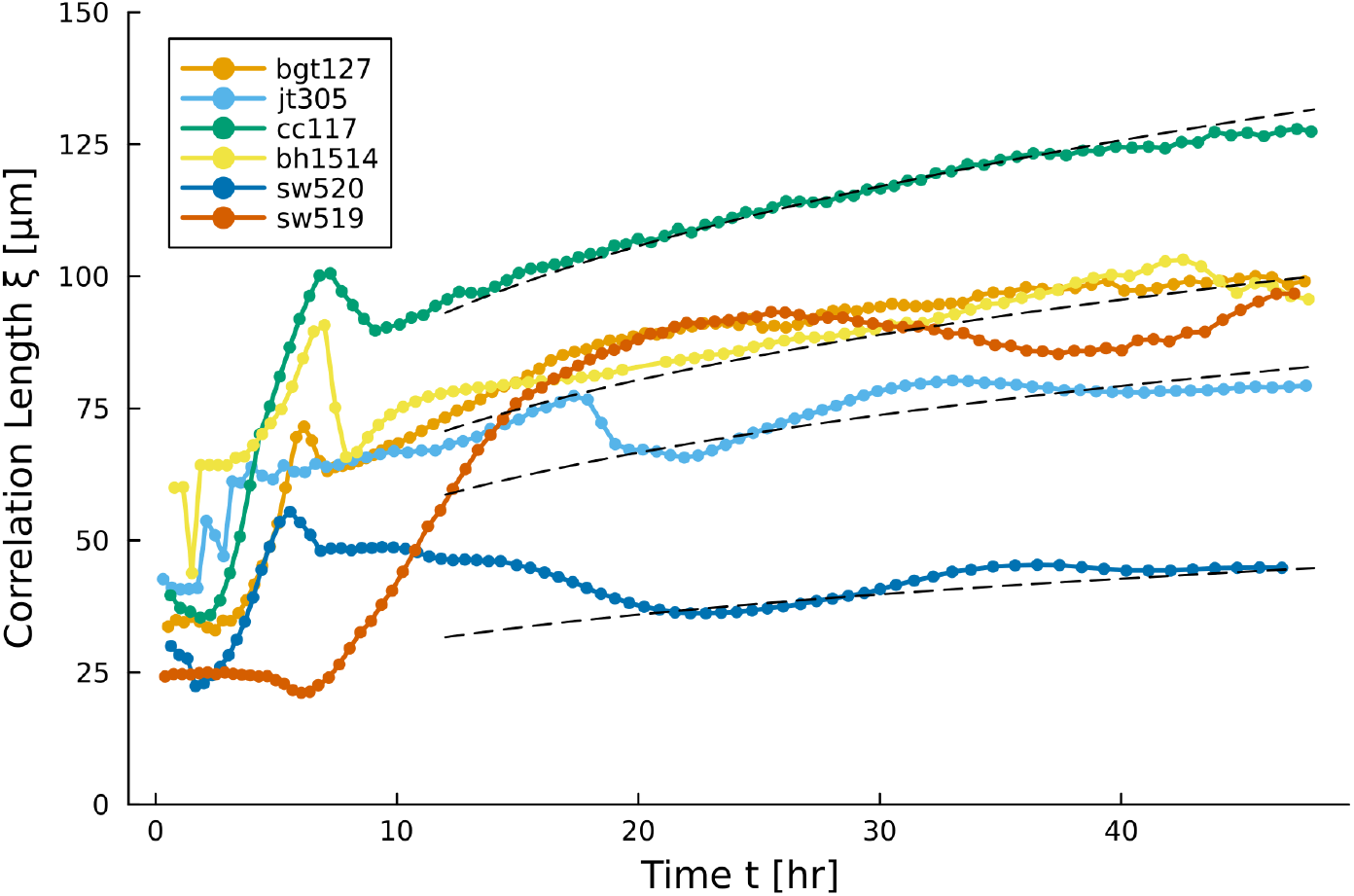
Correlation length scaling. Correlation length *ξ* evolution during the 48 hours of development. Dashed black lines correspond to *ξ* = *k* ^*¼*^, for 4 values of *k*, highlighting that the scaling behavior holds.

**Figure S5.**
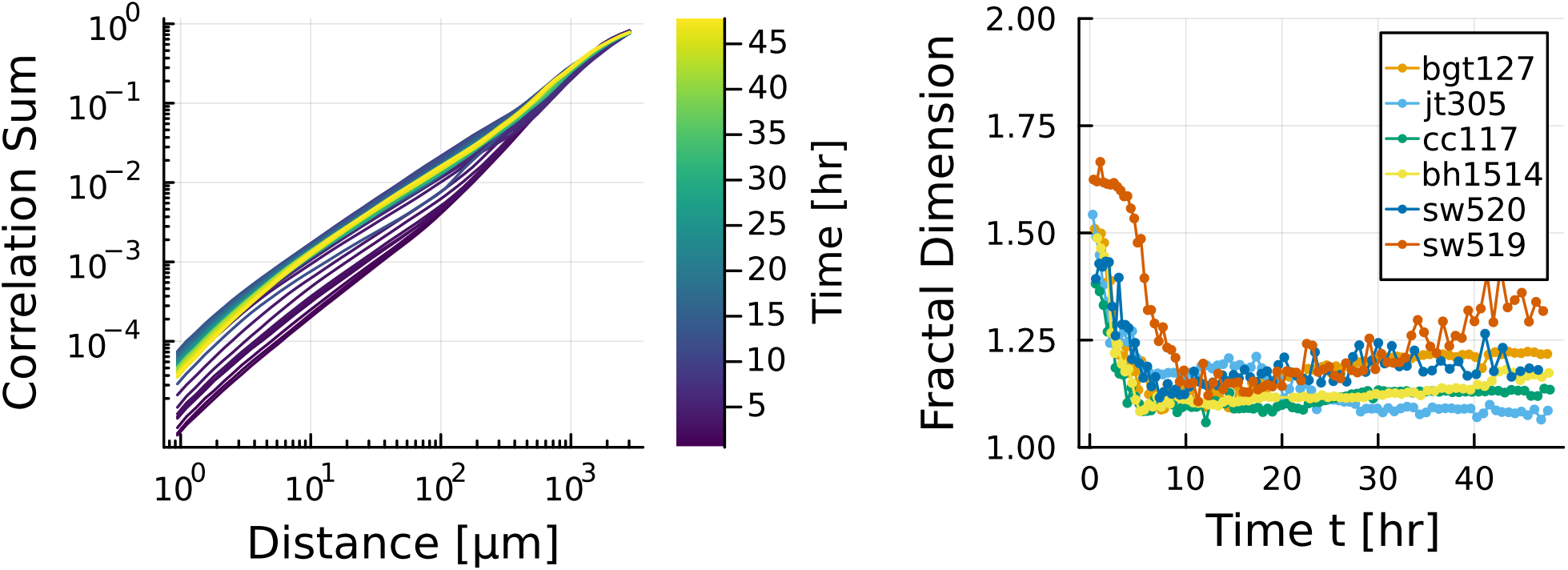
Fractal Dimension in biofilm-air interfaces. (Left) Correlation sum for a *Klebsiella pneumoniae* colony in relation to the distance *r*. The fractal dimension sets a scaling in the form goes as *C*(*r*) ∼ *rD*. The linear region in log-log is identified automatically, but correlates with the *r* = [10^0^, 10^2^] range. (Right) Fractal Dimension *D* behavior over time for species. The “freezing” of the topography corresponds to when *D* is not decreasing rapidly.

**Figure S6.**
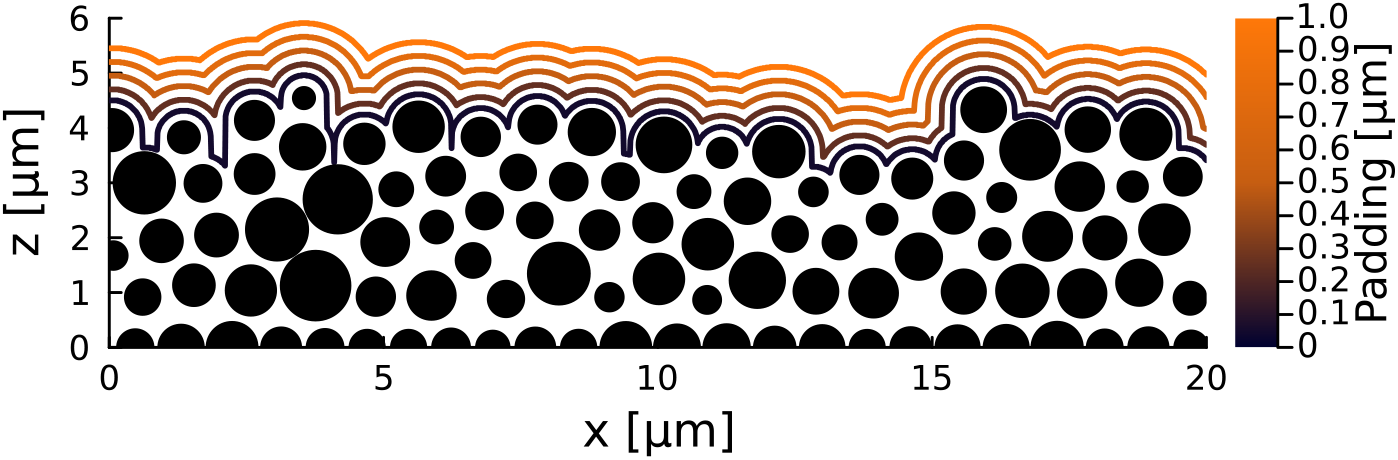
Topographies in an agent based model for biofilms. Different padding distances for a simulated colony. At low padding values, the interface has large dips when cells are apart. At high padding distances, the interface is too homogeneous and does not reflect the cell configurations accurately.

## References

1. Andrew Zangwill. Physics at surfaces. Cambridge university press, 1988.

2. Clarence A Miller and Partho Neogi. Interfacial phenomena: equilibrium and dynamic effects. CRC Press, 2007.

3. Timothy Halpin-Healy and Yi-Cheng Zhang. Kinetic roughening phenomena, stochastic growth, directed polymers and all that. aspects of multidisciplinary statistical mechanics. Physics reports, 254(4-6):215–414, 1995. doi:10.1016/0370-1573(94)00087-J.

4. A-L Barabási and Harry Eugene Stanley. Fractal concepts in surface growth. Cambridge university press, 1995.

5. Luanne Hall-Stoodley, J William Costerton, and Paul Stoodley. Bacterial biofilms: from the natural environment to infectious diseases. Nature Reviews Microbiology, 2(2):95–108, February 2004. doi:10.1038/nrmicro821.

6. Hans-Curt Flemming, Jost Wingender, Ulrich Szewzyk, Peter Steinberg, Scott A Rice, and Staffan Kjelleberg. Biofilms: an emergent form of bacterial life. Nature Reviews Microbiology, 14(9):563–575, August 2016. doi:10.1038/nrmicro.2016.94.

7. Philip S Stewart and Michael J Franklin. Physiological heterogeneity in biofilms. Nature Reviews Microbiology, 6(3):199–210, 2008. doi:10.1038/nrmicro1838.

8. Jeanyoung Jo, Alexa Price-Whelan, and Lars EP Dietrich. Gradients and consequences of heterogeneity in biofilms. Nature Reviews Microbiology, 20(10):593–607, 2022. doi:10.1038/s41579-022-00692-2.

9. Tohey Matsuyama, Masakazu Sogawa, and Yoji Nakagawa. Fractal spreading growth of serratia marcescens which produces surface active exolipids. FEMS microbiology letters, 61(3):243–246, 1989.

10. Hiroshi Fujikawa and Mitsugu Matsushita. Fractal growth of bacillus subtilis on agar plates. Journal of the physical society of japan, 58(11):3875– 3878, 1989. doi:10.1016/0378-1097(89)90204-8.

11. Juan A Bonachela, Carey D Nadell, João B Xavier, and Simon A Levin. Universality in bacterial colonies. Journal of Statistical Physics, 144:303– 315, 2011. doi:10.1007/s10955-011-0179-x.

12. Julien Dervaux, Juan Carmelo Magniez, and Albert Libchaber. On growth and form of Bacillus subtilis biofilms. Interface Focus, 4(6), 2014. doi: 10.1098/rsfs.2013.0051.

13. Alejandro Martínez-Calvo, Tapomoy Bhattacharjee, R. Kōnane Bay, Hao Nghi Luu, Anna M. Hancock, Ned S. Wingreen, and Sujit S. Datta. Morphological instability and roughening of growing 3D bacterial colonies. Proceedings of the National Academy of Sciences, 119(43):e2208019119, October 2022. doi:10.1073/pnas.2208019119.

14. Pablo Bravo, Siu Lung Ng, Kathryn A. MacGillivray, Brian K. Hammer, and Peter J. Yunker. Vertical growth dynamics of biofilms. Proceedings of the National Academy of Sciences, 120(11):e2214211120, March 2023. doi:10.1073/pnas.2214211120.

15. Pablo Bravo, Siu Liu Ng, Kathryn A. MacGillivray, Brian K. Hammer, and Peter J. Yunker. Radial topographies of biofilm colonies, 2023. doi:10.5061/dryad.pg4f4qrsw.

16. Kun Hu, Plamen Ch Ivanov, Zhi Chen, Pedro Carpena, and H Eugene Stanley. Effect of trends on detrended fluctuation analysis. Physical Review E, 64(1):011114, 2001. doi:10.1103/PhysRevE.64.011114.

17. Benoit B Mandelbrot and Benoit B Mandelbrot. The fractal geometry of nature, volume 1. WH freeman New York, 1982.

18. Tilmann Gneiting and Martin Schlather. Stochastic models that separate fractal dimension and the hurst effect. SIAM review, 46(2):269–282, 2004. doi:10.1137/S0036144501394387.

19. Jun-ichi Wakita, Hiroto Itoh, Tohey Matsuyama, and Mitsugu Matsushita. Self-affinity for the growing interface of bacterial colonies. Journal of the Physical Society of Japan, 66(1):67–72, 1997. doi:10.1143/JPSJ.66.67.

20. Peter Grassberger and Itamar Procaccia. Characterization of strange attractors. Physical review letters, 50(5):346, 1983. doi:10.1103/PhysRevLett.50.346.

21. George Datseris, Inga Kottlarz, Anton P Braun, and Ulrich Parlitz. Estimating fractal dimensions: A comparative review and open source implementations. Chaos: An Interdisciplinary Journal of Nonlinear Science, 33(10), 2023. doi:10.1063/5.0160394.

22. Arthur A Evans, Elliott Cheung, Kendra D Nyberg, and Amy C Rowat. Wrinkling of milk skin is mediated by evaporation. Soft Matter, 13(5):1056– 1062, 2017. doi:10.1039/c6sm02102f.

23. Stuart A Kauffman. The origins of order: Self-organization and selection in evolution. Oxford University Press, USA, 1993.

24. Per Bak. How nature works: the science of self-organized criticality. Springer Science & Business Media, 2013.

25. Mehran Kardar, Giorgio Parisi, and Yi-Cheng Zhang. Dynamic scaling of growing interfaces. Physical Review Letters, 56(9):889, 1986. doi: 10.1103/PhysRevLett.56.889.

26. Ivan Corwin. The kardar–parisi–zhang equation and universality class. Random matrices: Theory and applications, 1(01):1130001, 2012. doi:10.1142/S2010326311300014.

27. Paul Meakin. Fractals, scaling and growth far from equilibrium, volume 5. Cambridge university press, 1998.

28. Cristian Picioreanu, Mark CM Van Loosdrecht, and Joseph J Heijnen. Mathematical modeling of biofilm structure with a hybrid differential-discrete cellular automaton approach. Biotechnology and bioengineering, 58(1):101–116, 1998.

29. FDC Farrell, Oskar Hallatschek, D Marenduzzo, and B Waclaw. Mechanically driven growth of quasi-two-dimensional microbial colonies. Physical review letters, 111(16):168101, 2013. doi:10.1103/PhysRevLett.111.168101.

30. Ellen Young, Gavin Melaugh, and Rosalind J Allen. Active layer dynamics drives a transition to biofilm fingering. npj Biofilms and Microbiomes, 9(1):17, 2023. doi:10.1038/s41522-023-00380-w.

31. Isaac Klapper and Jack Dockery. Finger formation in biofilm layers. SIAM Journal on Applied Mathematics, 62(3):853–869, 2002. doi:10.1137/S0036139900371709.

